# Phage-Encoded TelN Inhibits Mre11-Rad50 to Protect Hairpin Telomeres

**DOI:** 10.1101/2025.06.15.659773

**Authors:** Maya Houmel, Nicolas Pellaton, Anna Anchimiuk, Stephan Gruber

**Affiliations:** Department of Fundamental Microbiology (DMF), Faculty of Biology and Medicine (FBM), University of Lausanne, 1015 Lausanne, Switzerland

**Keywords:** Rad50, Mre11, TelN, N15 phage, SMC-like, DNA end protection, telomeres, linear DNA, hairpin ends, DNA repair, phage-host interaction, telomere resolvase, broken ends, nuclease

## Abstract

The ends of linear chromosomes require protection from host repair machinery that otherwise will mistake them for damaged DNA. The *E. coli* bacteriophage N15 harbors a linear genome with covalently closed hairpin ends formed by the phage-encoded telomere resolvase TelN. The double-strand break repair complex Mre11-Rad50 (MR) specifically targets DNA termini, posing a direct threat to N15 genome integrity, yet how hairpin telomeres evade host nuclease degradation in bacteria remains unknown. Here, we demonstrate that TelN is essential and sufficient to protect hairpin telomeres from MR processing in *E. coli*. Using a combination of genetic and biochemical approaches, we show that this protective function requires both TelN sequence-specific DNA binding and species-specific protein- protein interactions. Notably, we found that protection is independent of TelN’s resolution activity and does not require the C-terminal domain of TelN. Our findings reveal a potentially broad mechanism of telomere protection, providing insights into a conserved regulation of MR activity at chromosome ends across the tree of life.

## Introduction

Maintaining genomic termini presents two fundamental challenges for organisms with linear chromosomes across all domains of life. First, semi-conservative replication fails to fully duplicate chromosome termini, known as the end replication problem, resulting in the need for dedicated enzymes such as telomerase to prevent progressive sequence loss. Second, linear chromosome ends must be protected from inappropriate processing by DNA repair machinery, referred to as the end protection problem. To address these fundamental challenges, eukaryotic cells have evolved specialized nucleoprotein structures called telomeres, to prevent failures in end replication and protection leading to nucleolytic degradation, chromosome end-to-end fusion, and ultimately cell cycle arrest or cell death (de Lange, 2009; Lazzerini-Denchi and Sfeir, 2016).

While most bacterial and archaeal chromosomes are circular, lacking DNA ends, some prokaryotic organisms harbor linear replicons, requiring protective mechanisms to preserve genome integrity. Bacterial telomeres fall into two distinct categories (Chaconas and Kobryn, 2010; Volff and Altenbuchner, 2000). The first type comprises terminal inverted repeat (TIR) sequences bound by terminal proteins, as found, for instance, in *Streptomyces* (Bao and Cohen, 2001) and in the *Bacillus subtilis* phage phi29 (Yoshikawa and Ito, 1981). The other category consists of covalently closed hairpin DNA ends, identified in diverse organisms, including the Lyme spirochete *Borrelia burgdorferi* (Fraser et al., 1997), the plant pathogen *Agrobacterium tumefaciens* (Goodner et al., 2001) as well as in various bacteriophages such as *Klebsiella oxytoca* phage phiKO2 (Stoppel et al., 1995) and *E. coli* phage N15 (Rybchin and Svarchevsky, 1999).

The mechanism of hairpin telomere replication has been extensively studied in the *E. coli* bacteriophage N15. The N15 prophage does not integrate into the host bacterial genome but exists as a linear DNA molecule with covalently closed hairpin ends. During N15 replication, the telomere resolvase TelN linearizes a dimeric circular replication intermediate by cleaving at two specific telomeric junction sites and sealing the ends, generating two monomeric linear copies of the chromosome with covalently closed hairpin telomeres (Deneke et al., 2000; Ravin, 2003; Ravin et al., 2001). TelN cleaving- joining activity requires its cognate *ntelRL* sequence, a 56-bp palindromic core essential for TelN binding and catalytic activity with additional repeats found in the flanking DNA sequences (Deneke et al., 2002). Following site-specific resolution, TelN generates a linear DNA molecule with covalently closed hairpin ends designated *ntelL* (left end) and *ntelR* (right end), obtained from processing of *ntelRR* and *ntelLL* sites produced by DNA replication (Ravin, 2015). Notably, the N15 TelN-*ntelRL* system was used to linearize the 4.6 Mb *E. coli* circular genome, resulting in viable cells with either a single linear chromosome (Cui et al., 2007) or with two complementary linear fragments (Liang et al., 2013), harboring covalently closed hairpin telomeres. These cells exhibited normal physiology, aside from the dispensability of the chromosome dimer resolution pathway (*dif*, XerCD), as expected for linear chromosomes.

While TelN effectively addresses the end replication problem through hairpin telomere creation and maintenance, how these linear structures evade processing by host nucleases is unknown. A significant threat to hairpin telomeres is the conserved Mre11-Rad50 (MR) complex, a key component of cellular double-strand break DNA repair machinery (Hopfner, 2023; Rojowska et al., 2014). The MR complex (also known as SbcCD in bacteria, including *E. coli*) comprises a Rad50 ATPase dimer associated with an Mre11 nuclease dimer, with an additional regulatory factor in eukaryotes (Nbs1 in mammals or Xrs2 in yeast) forming the MRN/MRX complex. MR specifically discriminates and processes linear DNA through a conserved mechanism in which Rad50 senses linear DNA ends, triggering ATP-dependent conformational changes that activate Mre11 nuclease activity (Käshammer et al., 2019); (Figure S1A). This enables both strand-specific endonucleolytic and 3’-5’ exonucleolytic cleavage at diverse DNA end structures, including free termini, protein-blocked ends, and hairpins (Connelly et al., 1998, 1999, 2003; Eykelenboom et al., 2008; Gut et al., 2022; Paull, 2018; Saathoff et al., 2018).

The MR complex, though essential for DNA repair, must be tightly regulated at telomeres. In eukaryotes, multiple parallel mechanisms protect chromosomal termini. Mammalian telomeres form compact T-loop structures through the shelterin complex, sequestering DNA ends (Van Ly et al., 2018; Smith et al., 2020), while in budding yeast, multiple Rap1 proteins sterically bind and cap telomeric regions, presumably stiffening DNA (Le Bihan et al., 2013). Simultaneously, three evolutionarily distinct peptide motifs are known to inhibit MRN/MRX directly. The iDDR motif in mammalian shelterin protein TRF2, MIN in telomere-associated Taz1 (fission yeast), and BAT in telomere-associated Rif2 (budding yeast), all target the same β-sheet of RAD50 to block nucleolytic processing at chromosome ends (Bombarde et al., 2010; Fan et al., 2025; Khayat et al., 2021, 2024; Marsella et al., 2021; Myler et al., 2023; Okamoto et al., 2013; Roisné-Hamelin et al., 2021).

Similar protection strategies against MR are needed in prokaryotes with linear replicons but remain largely uncharacterized. Hairpin telomeres are efficiently processed by the MR complex (Saathoff et al., 2018), yet a protection mechanism from inadvertent degradation must be in place for N15 propagation. In this study, we discover a role for TelN in hairpin telomere protection from *E. coli* MR processing, which depends on both sequence-specific DNA binding and species-specific protein- protein interactions. This protective function is distinct from TelN’s well-known catalytic activity and independent of its C-terminal domain. Our data suggest that MR inhibition at telomeres may represent one of the earliest evolutionary solutions to the challenge of maintaining linear chromosomes, establishing fundamental principles of telomere protection conserved from bacteria to humans.

## Results

### Chromosome linearization is lethal in the presence of host DNA repair nuclease Mre11-Rad50 in *B. subtilis*

To assess whether bacterial chromosomes can be linearized across different species, we attempted to generate linear chromosomes in *B. subtilis* using the phage-encoded telomere resolvase TelN and its *ntelRL* core sequence extended by neighboring sequences from the N15 phage (Figure 1A), an approach previously shown to successfully linearize the *E. coli* chromosome (Cui et al., 2007). We therefore engineered a *B. subtilis* strain containing this *ntelRL* sequence. Despite multiple attempts, no clones carrying both *telN* and intact *ntelRL* could be obtained, with rare suppressor clones carrying various deletions or mutations in the *ntelRL* insertion site (Supplementary Figure S1B). We wondered whether the generated linear chromosomes might be unstable due to their targeting by host DNA repair nucleases. To test this, we repeated the experiment in a strain lacking a gene for Mre11-Rad50 (ΔMR_*ntelRL*). Strikingly, transformation with a xylose-inducible *telN* construct yielded significantly more colonies in the ΔMR_*ntelRL* strain compared to wild type, where only a few, small-sized colonies were observed (Figure 1B). These results suggest that MR actively prevents the formation or maintenance of linear chromosomes in *B. subtilis*, but not in *E. coli* (Cui et al., 2007).

**Figure 1.**
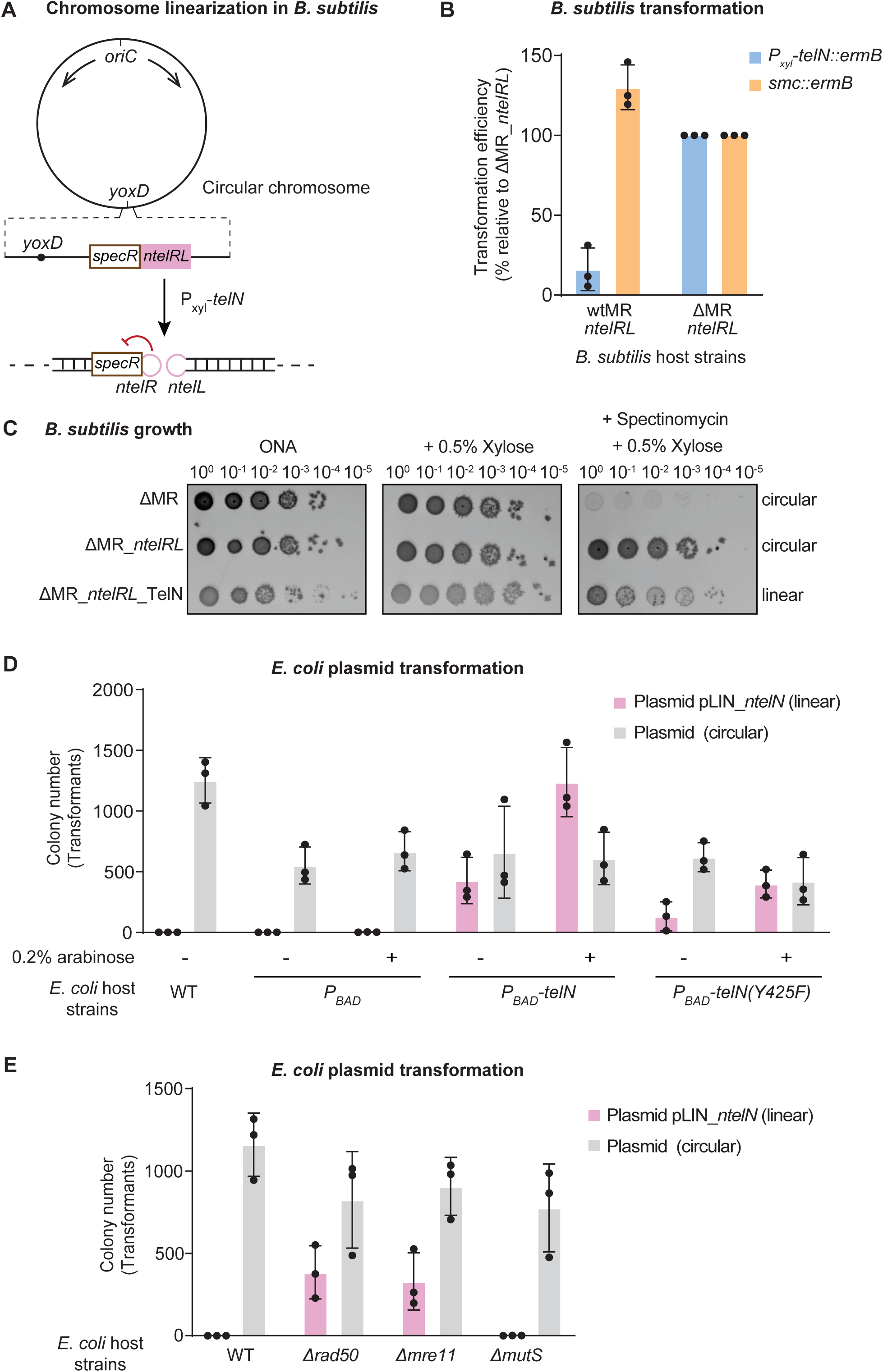
Antagonistic roles of Mre11-Rad50 and TelN in maintenance of linear DNA with hairpin **ends**. **(A)** The *ntelRL* sequence and spectinomycin resistance marker (*specR*) were integrated into the circular *B. subtilis* chromosome at the *yoxD* locus. Upon induction of TelN expression under the xylose promoter (*P_xyl_-telN*), TelN cleaves at the *ntelRL* site using its cleaving-joining activity, resulting in a linear *B. subtilis* chromosome with hairpin termini. The formation of these hairpin telomeres alters gene expression, resulting in reduced spectinomycin resistance (red arrow). **(B)** Counts of erythromycin-resistant colonies for the indicated *B. subtilis* backgrounds after transformation with TelN (*Pxyl*-*telN::ermB*) or Smc *(smc::ermB)* constructs. Both constructs are marked with erythromycin resistance cassettes (*ermB*), with TelN expression controlled by a xylose-inducible promoter (*P_xyl_*). Data are normalized to ΔMR_*ntelRL* (set as 100%). Means and standard deviations from three independent experiments are shown. Transformation with the *smc* control gene yielded similar colony numbers in both strains, confirming unimpaired natural competence in the wild-type background. **(C)** Serial spot dilutions (10°- 10^-5^) of *B. subtilis* ΔMR strains expected to carry either circular or linear replicons: ΔMR circular, ΔMR_*ntelRL* circular, and ΔMR_*ntelRL*_TelN linear. Cells were spotted on nutrient agar (ONA) plates supplemented as indicated. Growth was assessed under three conditions: ONA, ONA + 0.5% xylose, and ONA + spectinomycin 50 μg/mL + 0.5% xylose. **(D)** Counts of chloramphenicol-resistant colonies for the indicated *E. coli* backgrounds after transformation with either a linear plasmid, with *ntel* hairpin ends (*ntelL* and *ntelR*) and its own copy of the *telN* gene, denoted as pLIN_*ntelN*, or a control circular plasmid. Strains were grown with (+) or without (-) 0.2% arabinose. WT: wild-type strain, no chromosomal integration, P_BAD_: WT strain with chromosomal integration of arabinose-inducible promoter alone, *P_BAD_-telN*: WT strain with chromosomal integration of arabinose-inducible *telN* gene, *P_BAD_-telN(Y425F)*: WT strain with chromosomal integration of arabinose-inducible *telN(Y425F)*. Means and standard deviations from three independent experiments are shown. **(E)** Counts of chloramphenicol-resistant colonies for the indicated *E. coli* backgrounds after transformation with either the linear plasmid pLIN_*ntelN* or a control circular plasmid. Means and standard deviations from three independent experiments are shown.

To confirm and further characterize linear chromosome formation in *B. subtilis*, we first used antibiotic sensitivity as a proxy for genomic linearization, employing a spectinomycin resistance gene inserted at *ntelRL,* which becomes sensitized to the DNA hairpin formation likely due to loss of DNA super helicity near the end (Figure 1A). We observed two distinctive growth phenotypes in *B. subtilis* strains with linearized chromosomes. First, strains with a linear chromosome displayed impaired growth and the frequent appearance of suppressors in the absence of xylose, likely due to a requirement of TelN for telomere resolution (Figure 1C). Second, these strains showed reduced spectinomycin resistance, consistent with lowered gene expression at the DNA end, likely due to loss of DNA underwinding (Figure 1C). Moreover, we performed Southern blotting to confirm the linearization of *ntelRL* DNA on the *B. subtilis* chromosome (Supplementary Figure S1C and S1D). Taken together, these data provide strong evidence for successful chromosome linearization in *B. subtilis* strains lacking Mre11-Rad50 but not in wild type.

### TelN is required for protection of hairpin telomeres in *E. coli*

We next investigated how *E. coli* stably maintains linear DNA molecules despite the presence of MR. As a simple model system, we chose the phage N15-derived linear plasmid (previously commercialized under the name pJAZZ), here denoted as pLIN_*ntelN* (Godiska et al., 2010). pLIN_*ntelN* carries its own copy of the *telN* gene necessary for its replication as well as a chloramphenicol-resistant marker for selection of transformants (Supplementary Figure S1E). We first assessed the transformation efficiency of *E. coli* host cells containing or lacking an additional *telN* gene. Briefly, *telN* was placed under an arabinose-inducible promoter (P_BAD_) and integrated into the *E. coli* K12 chromosome near the *glmS* locus by mini-Tn7 transposon-based insertion (Bao et al., 1991). Strikingly, in the absence of chromosome-encoded *telN*, transformation of linear DNA was severely impaired compared to a circular control plasmid (Figure 1D). However, transformation of linear and circular DNA was similarly efficient when TelN was already present in host cells prior to transformation (Figure 1D). The presence of host- encoded *telN* is therefore needed for efficient uptake or maintenance of the linear plasmid with hairpin telomeres, presumably until plasmid-encoded TelN accumulates to sufficient levels.

Based on our previous findings in *B. subtilis* (Figure 1B), we suspected that the *E. coli* MR may be responsible for the low transformation efficiency of pLIN_*ntelN* in the absence of host-encoded *telN*. Thus, we transformed both linear and circular plasmids into *E. coli* strains deficient in the *rad50* or *mre11* genes. We observed that the absence of the *rad50* or *mre11* genes resulted in increased transformation efficiency of pLIN_*ntelN* to the level obtained by transformation with a circular plasmid, even in the absence of host-encoded *telN* (Figure 1E). This is not the case when deleting an unrelated gene, such as *mutS* in the host strain (Figure 1E). These results demonstrate that MR is indeed responsible for the low transformation efficiency of pLIN_*ntelN* and suggest that TelN protects linear DNA from MR-mediated degradation in *E. coli*.

### Direct inhibition of Mre11-Rad50 by TelN *in vitro*

To determine whether TelN directly inhibits Mre11-Rad50 nuclease activity, we developed an *in vitro* DNA degradation assay with a fluorescently labeled linear DNA substrate with a hairpin end. This substrate was generated by PCR amplification of the *ntelRL* sequence, along with adjacent N15 phage genomic regions, using a 5‘ biotinylated primer labeled with fluorescein (Figure 2A). TelN-mediated resolution of the *ntelRL* PCR product yielded fragments with one hairpin end. The fluorescently labeled hairpin substrate was immobilized on streptavidin-coated magnetic beads, using the physical size of the beads to sterically protect the biotin-labelled non-hairpin end and thus allowing MR to process the DNA from only one side, mimicking its suspected activity at unprotected telomeres (Figure 2A).

**Figure 2.**
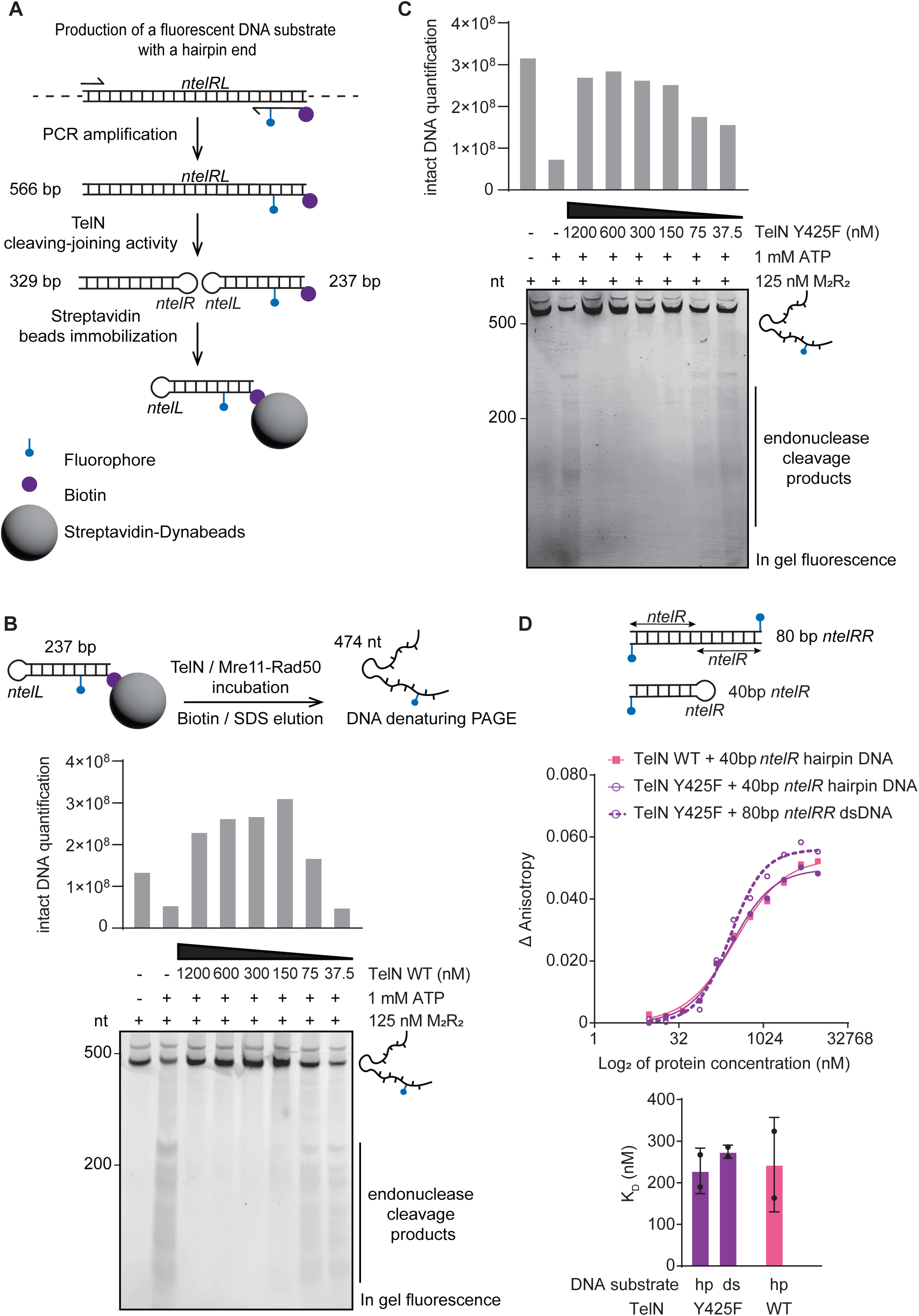
TelN directly inhibits Mre11-Rad50 nuclease activity independently of its catalytic function. **(A)** Schematic representation of the generation of a fluorescent hairpin DNA substrate for *in vitro* protection assays. The *nteIRL+* sequence (*ntelRL* cognate sequence with additional neighboring sequences from N15 phage) was PCR-amplified using a 3’-fluorescein/biotin-labeled reverse primer to generate a 566 bp product. TelN cleaving-joining activity produced two linear fragments with hairpin ends of 329 bp and 237 bp. The fluorescently labeled hairpin was immobilized on streptavidin-coated magnetic beads via biotin for further analysis. **(B-C)** *In vitro* protection assays using **(B)** TelN WT or **(C)** TelN Y425F with Mre11-Rad50. 37 nM of the immobilized *ntelL* hairpin substrate was incubated with 125 nM M_2_R_2_ and decreasing concentrations of the respective TelN variant. Products were analyzed by denaturing PAGE after biotin/SDS elution. Under denaturing conditions, the hairpin DNA (237 bp) unfolds and runs at twice its size (474 bp). Lower panels: representative in-gel fluorescence analysis showing DNA products. Upper panels: quantification of intact DNA substrates. **(D)** DNA binding analysis by fluorescence anisotropy. Upper panel: binding curves of TelN WT with 40 bp *ntelR* hairpin DNA and TelN Y425F with 40 bp hairpin DNA or 80 bp *ntelRR* dsDNA. Lower panel: calculated dissociation constants (KD) for different DNA substrates and TelN variants. Error bars represent standard deviations from two independent experiments.

We added purified MR and TelN proteins (Supplementary Figures S2A and S2B) to the bead- immobilized hairpin substrate and observed that in the absence of TelN, MR efficiently degraded the DNA in the presence of ATP (Figure 2B). Interestingly, the addition of TelN inhibited MR nuclease activity in a dose-dependent manner (Figure 2B). This demonstrates that TelN is sufficient to protect hairpin telomeres from *E. coli* MR degradation without requiring additional cellular factors.

### Hairpin telomere protection is independent of TelN’s ability to resolve hairpin telomeres

Given that TelN is primarily characterized as a telomere resolvase, we next investigated whether its catalytic activity was required for hairpin telomere protection. We thus generated a catalytically inactive TelN Y425F mutant using site-directed mutagenesis (Deneke et al., 2000) (Supplementary Figure S2B). As expected, purified TelN Y425F showed no detectable DNA cleavage activity on the *ntelRL* target sequence even at high protein concentrations, while TelN WT efficiently processed the substrate (Supplementary Figures S2C and S2D). Strikingly, we observed that TelN Y425F inhibited MR-mediated hairpin degradation as effectively as TelN WT in our *in vitro* protection assay (Figure 2C). This suggests that TelN’s resolvase and protection activities can be separated. To confirm these observations in a cellular context, we expressed TelN Y425F from the *E. coli* chromosome and assessed its impact on telomere protection. Interestingly, TelN Y425F improved the transformation efficiency of linear plasmid with hairpin ends similar to WT (Figure 1D). This result further supports the notion that TelN’s protective function is mechanistically distinct from its telomere resolution activity.

### DNA end binding by TelN

To understand how TelN mediates protection, we first assessed its DNA binding properties by fluorescence anisotropy. We designed fluorescein-labeled DNA substrates containing the *ntelR* sequence in two conformations: a 40-bp *ntelR* hairpin conformation and an 80-bp linear *ntelRR* duplex (Supplementary Figure S2E). TelN bound to all specific substrates with moderate affinity (dissociation constant (K_D_) in the ∼200-300 nM range), regardless of the DNA structure or length (Figure 2D). Due to the catalytic activity of TelN, its binding to the linear 80 bp *ntelRR* dsDNA substrate could not be reliably measured, as cleavage would have likely occurred upon DNA recognition. As expected, no significant differences in binding affinity were observed between TelN WT and TelN Y425F on the 40 bp *ntelR* hairpin substrate, suggesting that the catalytic activity of TelN does not influence its DNA binding properties to single binding sites (Figure 2D). Surprisingly, the binding affinity of TelN Y425F remained comparable between the hairpin with a single binding site and dsDNA substrates with two binding sites, indicating a lack of strong cooperativity in DNA binding and implying that two TelN proteins bind largely individually to the two binding sites (Figure 2D). We conclude that TelN specifically recognizes its target sequence with moderate affinity. Our findings suggest that TelN binds to the educt and the product of the enzymatic reaction with similar affinity, implying that it remains bound to the target DNA after telomere resolution, unlike typical enzymes.

### The C-terminal domain of TelN is dispensable for hairpin telomere protection

The C-terminal domain of telomere resolvases is less well conserved across species, and its functions remain unknown (Supplementary Figure S3A). To investigate this domain’s role in TelN’s protective function, we generated two truncation variants lacking the C-terminal domain: Δ540-631 and Δ582- 631 (Figure 3A). We integrated these mutants into the *E. coli* chromosome under arabinose-inducible control to assess their ability to protect linear DNA *in vivo*. TelN WT provided protection even without induction. The Δ540-631 and Δ582-631 variants also protected but required arabinose induction to restore the transformation efficiency of linear plasmids to wild-type levels (Figure 3B). We next tested whether these truncated variants could protect hairpin telomeres from MR degradation *in vitro.* Purified Δ540-631 and Δ582-631 proteins (Supplementary Figure S3B) inhibited MR nuclease activity comparably to wild-type TelN (Figure 3C, 3D), while retaining full catalytic activity in telomere resolution assays (Supplementary Figure S3C, S3D). These results demonstrate that the C-terminal domain is dispensable for telomere protection, suggesting that the core protection mechanism resides in the conserved region of the protein.

**Figure 3.**
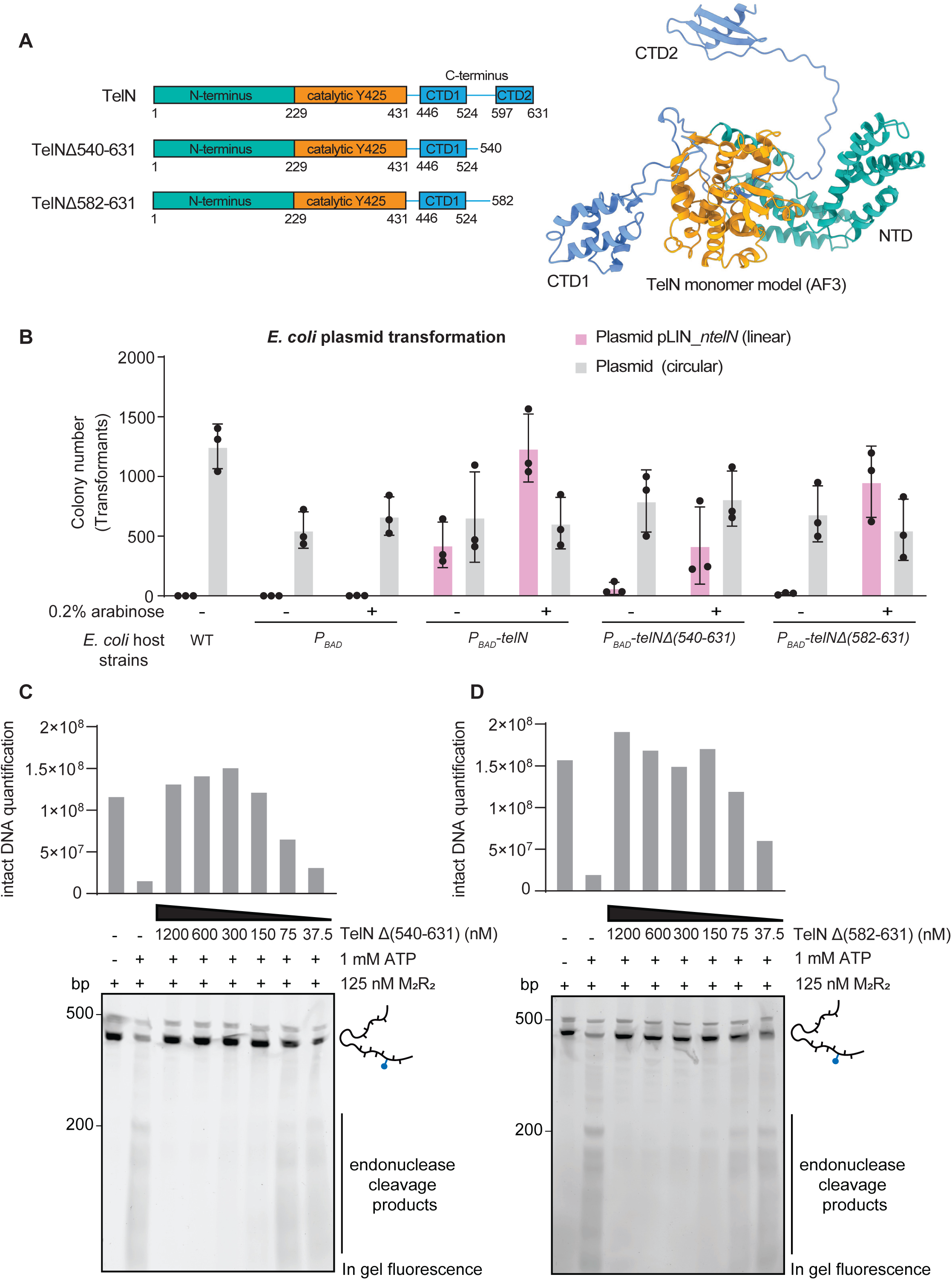
The C-terminal domain of TelN is dispensable for telomere protection. **(A)** Left panel: Domain organization of TelN showing N-terminus, catalytic Y425 region, and C-terminus with positions of truncations (Δ540-631 and Δ582-631) indicated. Right panel: Alphafold3 prediction of TelN monomer (pTM = 0.74) showing N-terminal domain in green (NTD), catalytic Y425 region in orange, and C-terminal domains in blue (CTD1 and CTD2). **(B)** Counts of chloramphenicol-resistant colonies for the indicated *E. coli* strains after transformation with either the linear plasmid pLIN_*ntelN* or a control circular plasmid. Strains were grown with (+) or without (-) 0.2% arabinose. WT: wild-type strain, no chromosomal integration, P_BAD_: WT strain with chromosomal integration of arabinose- inducible promoter alone, *P_BAD_-telN*: WT strain with chromosomal integration of arabinose-inducible *telN* gene, *P_BAD_*-*telN(Δ540-631)*: WT strain with chromosomal integration of arabinose-inducible *telN(Δ540-631)*, *P_BAD_-telN(Δ582-631)*: WT strain with chromosomal integration of arabinose-inducible *telN(Δ582-631)*. Means and standard deviations from three independent experiments are shown. **(C- D)** *In vitro* protection assays using **(C)** TelN(Δ540-631) or **(D)** TelN(Δ582-631) with Mre11-Rad50. 37 nM of the immobilized *ntelL* hairpin substrate (237 bp) was incubated with 125 nM M_2_R_2_ and decreasing concentrations of the respective TelN variant. Products were analyzed by denaturing PAGE after biotin/SDS elution. Under denaturing conditions, the hairpin DNA (237 bp) unfolds and runs at twice its size (474 bp). Lower panels: representative in-gel fluorescence analysis showing DNA products. Upper panels: quantification of intact DNA substrates.

### Sequence specificity of TelN-mediated protection is modulated by cellular context

To elucidate the molecular requirements for TelN-mediated protection, we tested whether binding of TelN to *ntelRL* was important for protection. We first tested *in vivo* whether TelN could protect non-cognate DNA sequences by comparing it with TelPY, a related telomere resolvase from *Yersinia enterocolitica* bacteriophage PY54 (Hertwig et al., 2003) that shares 43% sequence identity (Supplementary Figure S4A). We designed linear plasmids with two different hairpin ends for analysis (Figure 4A). Starting from pLIN_*ntelN*, we produced a circular form upon inactivation of the plasmid- encoded *telN* gene through an early stop codon and in the same cloning step, ligating *ntelR* and *ntelL* to recreate *ntelRL*. We then replaced the *ntelRL* recognition sequence by *pytelRR*, the recognition sequence of TelPY, which shares 45% sequence identity (Supplementary Figure S4B). This plasmid was transformed into a ΔMR *E. coli* strain expressing chromosomal *telPY*, yielding a linear plasmid with two *pytelR* hairpin ends (Figure 4A).

**Figure 4.**
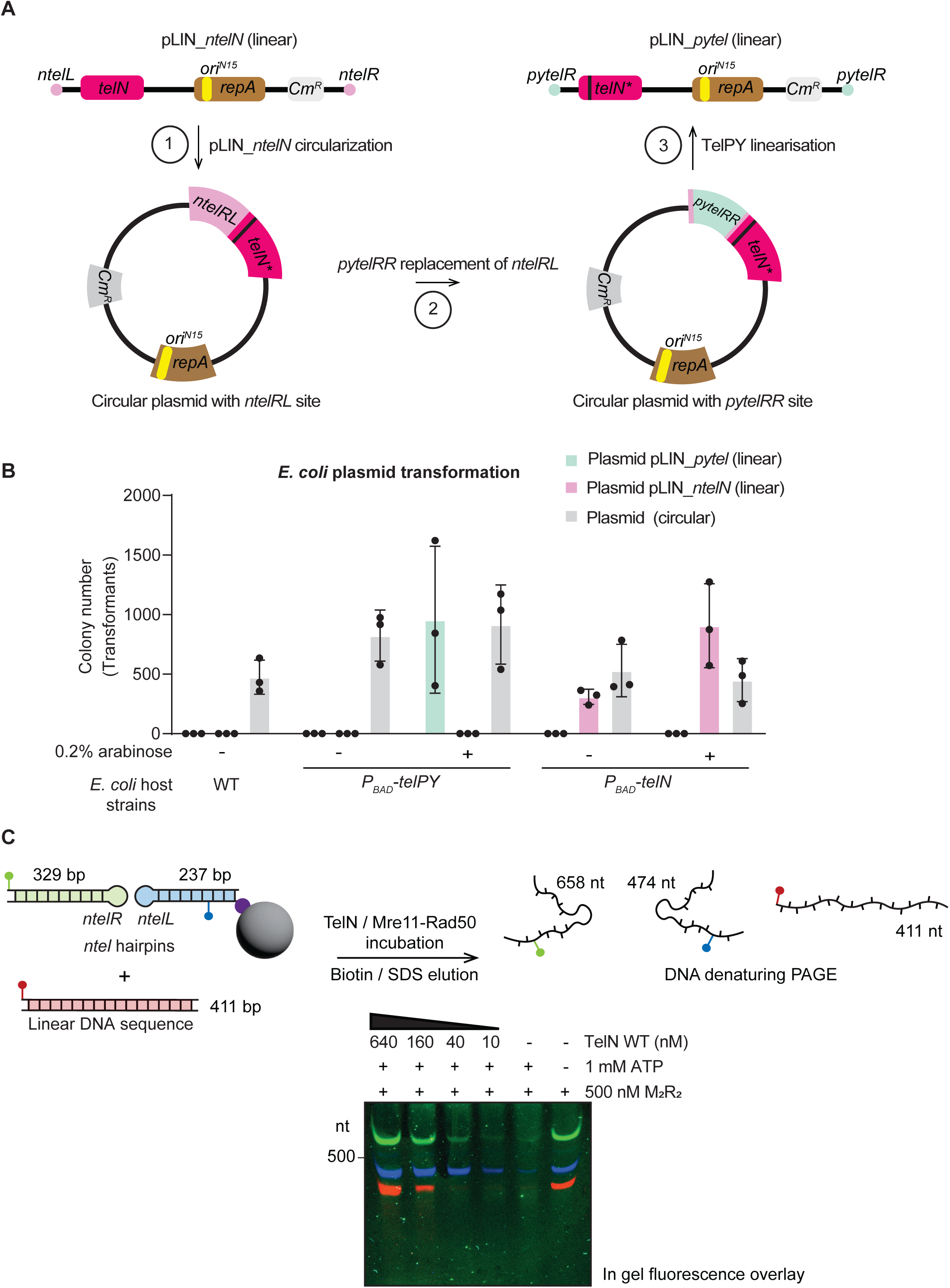
Sequence specificity of TelN-mediated protection differs between cellular and biochemical **contexts.** **(A)** Overview of the cloning procedure for generating linear plasmids with different hairpin ends. Left pathway: Linear plasmid with *ntel* hairpin ends, carrying its own copy of the *telN* gene, pLIN_*ntelN*. Step 1: Circularization to generate a circular plasmid containing the unresolved *ntelRL* sequence (*ntelRL* sequence with adjacent N15 phage genomic regions). Step 2: Replacement of the unresolved *ntelRL* with the unresolved *pytelRR* sequence. The *pytelRR* DNA substrate was constructed by duplicating *pytelR*, one half of the PY54 telomerase target site (*pytelRL*), along with neighboring sequences from the PY54 phage, in an inverted orientation to generate a palindromic structure. Step 3: TelPY-mediated linearization to generate a linear plasmid with *pytel* hairpin ends pLIN_*pytel*. Adapted from (Liu et al., 2022) **(B)** Counts of chloramphenicol-resistant colonies for the indicated *E. coli* strains after transformation with linear plasmids containing either *ntel* (pLIN_*ntelN*) or *pytel* (pLIN_*pytel*) hairpin ends, or a control circular plasmid. Strains were grown with (+) or without (-) 0.2% arabinose. WT: wild- type no chromosomal integration, *P_BAD_-telPY*: WT strain with chromosomal integration of arabinose- inducible *telPY*, *P_BAD_*_*telN*: WT strain with chromosomal integration of arabinose-inducible *telN*. Means and standard deviations from three independent experiments are shown. **(C)** Upper panel: Schematic representation of the *in vitro* protection assay. Three distinct fluorescently labeled DNA substrates were tested simultaneously: two fluorescently labeled *ntel* hairpin substrates of 329 bp and 237 bp, and one non-specific linear DNA fragment of 411 bp. The total DNA concentration was maintained at 30 nM, with equimolar amounts of each substrate (10 nM each). Lower panel: Representative in-gel fluorescence analysis showing DNA products after incubation with 500 nM M_2_R_2_ and decreasing concentrations of TelN. Fluorophore overlay image from Typhoon fluorescence scanner shows simultaneous visualization of all three denatured substrates (green/blue: *ntel* hairpin substrates appearing at 658 bp and 474 bp due to denaturation; red: non-specific linear DNA at 411 bp).

Using the two linear constructs, pLIN_*ntelN* with *ntel* and pLIN_*pytel* with *pytel* hairpin ends, we assessed protection specificity through plasmid transformation assays. Despite the sequence similarity between their respective binding sites, the proteins exclusively protected their cognate DNA sequence. Chromosomally encoded *telN* enabled efficient transformation of pLIN_*ntelN* even without arabinose induction, but failed to support transformation of pLIN_*pytel* (Figure 4B) even with induction. Similarly, TelPY showed similar sequence specificity by only protecting plasmids with its cognate *pytel* hairpin ends, though requiring arabinose induction for protection (Figure 4B). This strict specificity suggests that sequence-specific DNA binding is essential for protection, ruling out mechanisms that solely rely on a direct interaction between the telomere resolvase protein and the MR complex.

To gain more insights into this binding specificity, we examined the protective capacity of TelN under defined biochemical conditions. We assessed TelN’s ability to protect three distinct DNA substrates *in vitro*: two fluorescently labeled *ntel* hairpins (*ntelL* and *ntelR*) and a non-specific fluorescent linear DNA fragment lacking *ntelRL* sequences (Figure 4C). Interestingly, at slightly elevated protein concentrations, TelN also protected non-specific DNA ends from MR degradation, with nearly complete protection of all substrates at 640 nM TelN (Figure 4C, Supplementary Figures S4C, S4D, and S4E). This discrepancy between *in vivo* specificity and broader substrate protection *in vitro* suggests that spatial organization likely modulates TelN’s protective function in cells. While TelN can inhibit MR activity on various DNA substrates when present at high concentrations in solution, the cellular environment, with its vast excess of non-specific DNA, likely constrains this activity through sequence-specific recruitment, ensuring protection is directed specifically to telomeric sequences.

### Species specificity of TelN-mediated protection

To further characterize the protection mechanism, we investigated whether TelN could inhibit MR complexes from different species. We compared TelN’s ability to inhibit nucleolytic processing by both *E. coli* MR and a eukaryotic counterpart, the MRX complex from *S. cerevisiae*. Despite their functional similarity, these complexes only share limited sequence homology. *E. coli* MR proteins share only 10- 12% sequence identity with their yeast counterparts (Supplementary Figures S5A and S5B). Using a fluorescently labeled hairpin substrate immobilized on streptavidin-coated magnetic beads, we compared TelN’s protective activity against purified *E. coli* MR versus its eukaryotic ortholog MRX. TelN effectively blocked *E. coli* MR nuclease activity, while showing no protection against yeast MRX degradation (Figure 5A, 5B). This species-specificity suggests that TelN makes specific contacts with the bacterial MR complex, pointing toward a protection mechanism involving direct protein-protein interactions with the *E. coli* complex, thus providing an explanation for the lack of protection observed in *B. subtilis*.

**Figure 5.**
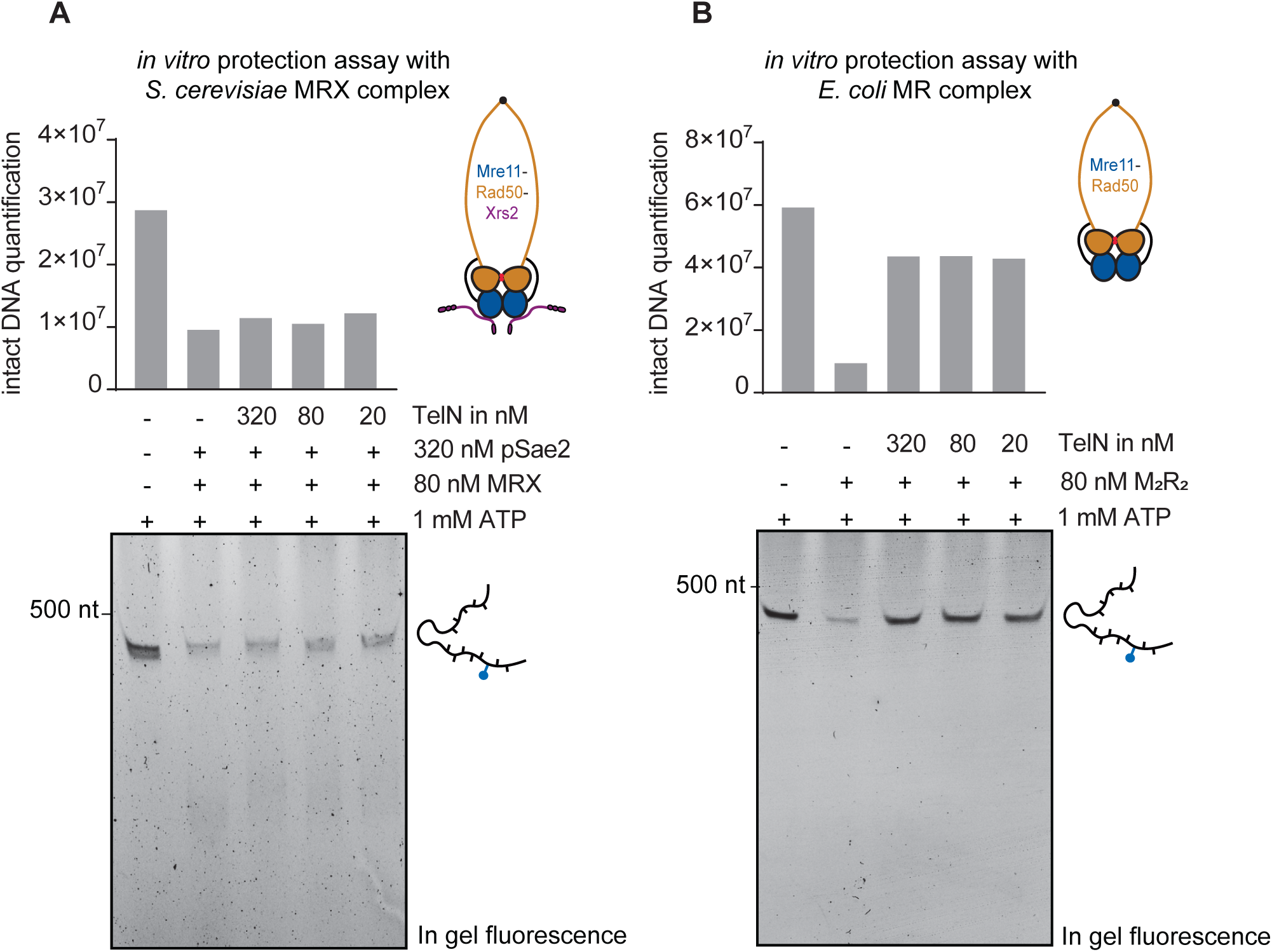
TelN-mediated protection exhibits species specificity. **(A)** *In vitro* protection assay testing TelN activity against *S. cerevisiae* MRX complex. Lower panel: representative in-gel fluorescence analysis showing DNA products after incubation with 320 nM pSae2 and 80 nM MRX and decreasing concentrations of TelN. Upper panel: quantification of intact DNA substrates. **(B)** *In vitro* protection assay testing TelN activity against *E. coli* MR complex. Lower panel: representative in-gel fluorescence analysis showing DNA products after incubation with 80 nM MR and decreasing concentrations of TelN. Upper panel: quantification of intact DNA substrates. The DNA substrate used for protection assays was a fluorescently labeled *ntelL* hairpin DNA (237 bp) immobilized on streptavidin-coated magnetic beads at 1 nM concentration. Under denaturing conditions, the hairpin DNA (237 bp) unfolds and runs at twice its size (474 bp).

## Discussion

Maintenance of chromosome ends and protection from cellular nucleases pose a fundamental challenge for organisms with linear chromosomes. Eukaryotes have evolved several independent mechanisms to protect their telomeres from degradation, prevent end-to-end chromosome fusions, and telomere erosion. In contrast, the mechanisms governing DNA end protection in bacteria with linear chromosomes remain poorly understood. This study demonstrates the role of phage-encoded TelN telomere resolvase in hairpin telomere protection. Using both *in vivo* and *in vitro* approaches, we show that TelN is essential and sufficient to protect hairpin telomeres from MR nuclease degradation in *E. coli* (Figures 1D and 2B). Notably, the hairpin resolvase catalytic activity of host-encoded TelN is not needed for efficient protection of linear DNA with hairpin ends (Figures 1D and 2C), revealing a clear separation of function.

### Molecular mechanisms underlying TelN-mediated end protection

Our findings suggest that TelN involves multiple coordinated steps to protect hairpin telomeres effectively. First, our data demonstrate a requirement for sequence-specific DNA recognition and binding, as TelN exclusively protects its cognate sequence *in vivo* (Figure 4B). However, TelN’s moderate binding affinity (Figure 2D) suggests that binding alone is not sufficient to confer effective protection. Second, the species-specific inhibition of *E. coli* MR but not yeast MRX indicates a direct protein-protein interaction between TelN and the bacterial MR complex (Figure 5). Our data suggests at least two steps or two features of the same mechanism acting in parallel. This dual requirement for both DNA binding and specific MR interaction mirrors strategies employed by eukaryotic telomere protection factors. For instance, in *S. cerevisiae*, the telomere-associated protein Rap1 recognizes and binds repeat sequences in tandem, presumably stiffening telomeric DNA (Le Bihan et al., 2013; Williams et al., 2010). Rap1 also recruits another telomere-associated protein, Rif2, which directly inhibits MRX through its BAT motif binding to Rad50 (Roisné-Hamelin et al., 2021).

The observed difference between TelN’s sequence-specific protection *in vivo* and its ability to protect non-specific DNA *in vitro* (Figures 4B and 4C) provides important insights into the spatial regulation of telomere protection. While TelN can inhibit MR activity on various DNA substrates at elevated concentrations in solution, its protective function requires precise binding to specific sequences in a cellular context. This mirrors the eukaryotic telomere protection factor Rif2, which inhibits MRX endonuclease activity *in vitro* without requiring DNA recruitment (Marsella et al., 2021), yet *in vivo*, it is spatially constrained through Rap1-dependent localization at telomeres (Roisné-Hamelin et al., 2021). This permits the distinction between chromosome ends that need to be protected from DNA damage sites that need to be repaired. Likely, TelN remains bound to newly generated DNA ends, thus ensuring protection of DNA ends even at low TelN concentrations.

Recent structural insights into MR conformational states during DNA end processing show that recognition of a DNA end induces a ring-to-rod transition of the coiled coils following ATP hydrolysis (Gut et al., 2022; Käshammer et al., 2019). The Mre11 dimer moves to the side, forming a nuclease proficient complex. TelN might hamper the conformational change of the MR complex, preventing ATP binding and hydrolysis, thus blocking the release of the nuclease and preventing DNA end processing. MR would be stuck in its ATP-bound state, unable to capture telomeric structures. Our *in vitro* assays support this hypothesis, demonstrating that TelN directly inhibits the nuclease activity of the MR complex (Figure 2B), thereby protecting DNA ends from degradation. A similar mechanism is proposed in eukaryotic systems, where, for instance, the iDDR motif in mammalian shelterin protein TRF2, MIN in telomere-associated Taz1 (fission yeast), and BAT in Rif2 (budding yeast) inhibit MRN/X-dependent DNA resection at telomeres by binding the same exposed β-sheet region of the RAD50 ATPase head (Fan et al., 2025; Khayat et al., 2024). The structural basis of potential TelN-MR interactions remains to be determined, and high-resolution structural studies using cryo-electron microscopy could reveal whether TelN induces conformational changes in the *E. coli* MR complex. Our attempts to detect direct interactions through co-immunoprecipitation between TelN and MR were unsuccessful, suggesting that their interaction might be transient or context-dependent. Additionally, AlphaFold-based structural predictions have not yet yielded informative models for this potential complex.

### Higher-order structure formation in DNA end protection?

Several observations suggest that TelN may induce structural changes in telomeric DNA for efficient protection. First, the natural target sequence of TelN is significantly larger (310 bp) than the *ntelRL* recognition site required for recombination (56 bp) (Deneke et al., 2002). This sequence includes additional R and L binding sites, implying the recruitment of multiple TelN proteins to a single DNA end. This might support the formation of larger nucleoprotein assemblies, with additional binding sites facilitating the creation of a more complex and stable structure. Second, our *in vitro* data demonstrate that even at low concentrations, TelN efficiently protects cognate DNA sequences from MR degradation (Figure 2B), which could potentially be explained by the formation of higher-order DNA structures beyond simple DNA binding. Third, structural analyses of related telomere resolvases, such as TelK from *Klebsiella oxytoca* phage phiKO2, bound to its minimal target DNA sequence (Aihara et al., 2007), reveal a basic nucleoprotein complex that would likely be insufficient for complete DNA end protection from MR. We therefore speculate that TelN contributes to the compaction of hairpin telomeres folding into a larger structure, such as the Telomere (T)-loops observed in mammals (Van Ly et al., 2018; Smith et al., 2020). TelN-mediated DNA looping can be envisaged as an architectural solution to protect hairpin telomeres. It would be an effective mechanism to mediate end protection by sequestering and masking the extreme chromosome termini.

### Evolutionary implications

Our findings provide fundamental insights into bacterial telomere biology and broaden our understanding of DNA end protection in prokaryotic systems. The dispensability of most of the TelN C- terminal domain for protection (Figure 3), which is highly variable among telomere resolvases, raises the possibility that other bacterial telomere resolvases might share similar protective functions. The parallels between bacterial and eukaryotic telomere protection suggest that conserved strategies have evolved to solve the end-protection problem. TelN-mediated protection of N15 hairpin ends is likely crucial for phage survival and propagation and a result of evolutionary adaptation, providing valuable insights into phage-host coevolution. In this regard, MR is acting as an anti-phage defense system with TelN providing counter-defense activity. How TelN evades counter-counter-defense mechanisms remains to be determined.

## Acknowledgements

We thank the Cejka lab for providing the MRX protein complex and the Sae2 protein. We thank Michael Taschner and Yan Li for help and advice on recombinant protein purification, Hon Wing Liu for feedback on figure schematics, and Florian Roisné-Hamelin for stimulating discussions and valuable input on data interpretation. We are grateful to the members of the Gruber lab for helpful feedback during manuscript preparation. This work was supported by SNSF project funding (320030-227915) to SG.

## Author contributions

MH and SG designed the assays. MH performed experiments with the following exceptions: SG isolated and characterized *ntelRL* suppressor mutations in wild-type *B. subtilis* (Supplementary Figure 1B). AA established chromosome linearization in *B. subtilis* MR mutants. Other experiments in *B. subtilis* (Figure 1B) including the Southern blot analysis of *B. subtilis* genomic DNA (Supplementary Figure 1D) were performed by NP. MH analyzed the data and wrote the first draft of the manuscript. SG supervised the project and obtained funding.

## Materials and Methods

### *B. subtilis* strain construction

All strains constructed in this work are derived from the 1A700 isolate. Natural competence was used to engineer strains at the *yoxD rtp proH*, *amyE*, *cgeD*, *sbcC, and sbcD* loci by allelic replacement, as described in (Diebold-Durand et al., 2019). Strains were selected on SMG-agar plates under appropriate antibiotic selection at 37°C. Genotypes were verified for single-colony isolates by PCR and Sanger sequencing (Microsynth) as required.

### Transformation assay in *B. subtilis*

Transformation assays were performed with naturally competent *B. subtilis* cells. To introduce genomic DNA, we used 60-80 ng of DNA from a strain carrying the appropriate chromosomal antibiotic resistance marker. Plates containing transformant colonies were imaged, and colonies were manually counted.

### Viability assessment by dilution spotting

Bacterial strains were initially cultured on Oxoid Nutrient Agar (ONA) plates supplemented with 0.5% xylose. Single colonies were subsequently inoculated into Luria-Bertani (LB) medium containing 0.5% xylose and cultured overnight at 30°C with continuous agitation. The overnight cultures were subjected to serial 1:10 dilutions, and bacterial density was estimated by measuring optical density at 600 nm (OD600). Fresh LB medium supplemented with 0.5% xylose was inoculated with the overnight culture to an initial OD600 of 0.001. These cultures were incubated at 37°C until reaching exponential phase (OD600 ≈ 0.3). For spotting assays, 200 μL of the exponential phase culture was serially diluted in a 96- well plate to achieve dilutions ranging from 10^−1^ to 10^−7^. Technical duplicates of 5 μL from each dilution were spotted onto ONA plates containing selective conditions. Colony growth was documented by imaging after 19 h of incubation at 37°C.

### *E. coli* strain construction

A mini-*Tn7* transposon carrying *araC* and different TelN constructs (TelN WT, TelN Y425F, TelN Δ540- 631, TelN Δ582-631) under the P_BAD_ promoter was integrated into a neutral *E. coli* chromosomal locus downstream of *glmS* by triparental mating, as previously described (Bao et al., 1991). This approach generated multiple *E. coli* strains, each expressing a distinct TelN variant.

### *E. coli* chemically competent cell preparation and transformation

To prepare chemically competent *E. coli* cells, a sterile flask containing the required volume of LB medium was inoculated with a few colonies from a freshly streaked plate and grown at 37°C with shaking until an OD600 of 0.4–0.6 was reached. The culture was chilled on ice for 15 min and harvested by centrifugation (5000 rpm, 10 min, 4°C). Cell pellets were gently resuspended in ice-cold TBF I buffer (30 mM potassium acetate, 100 mM RbCl, 10 mM CaCl3, 50 mM MnCl3*4H3O, 15% glycerol, pH 5.8), using a volume equivalent to 0.4× the original culture volume. After 5 min incubation on ice and subsequent centrifugation, pellets were resuspended in ice-cold TBF II buffer (10 mM MOPS, 10 mM RbCl, 75 mM CaCl3, 15% glycerol, pH 6.5) at 0.02× the original culture volume. Following 15 min incubation on ice, the suspension was aliquoted (100 μL), flash-frozen in liquid nitrogen, and stored at –70°C. Where indicated, 0.2% arabinose was added to the culture medium.

For transformation experiments, 1 ng of DNA (either linear plasmids with covalently closed hairpin ends or circular plasmids) was added to 100 μL chemically competent cell aliquots. After heat shock treatment at 42°C for 1 min, cells were recovered in LB medium for 1 h at 37°C. Transformed cells were then plated on LB agar supplemented with chloramphenicol and, where indicated, 0.02% arabinose. Plates were incubated overnight at 37°C. Colony counts were determined using ImageJ software, and all experiments were performed in triplicate.

### DNA substrate preparation and fluorescent anisotropy assay

Two DNA substrates were used for fluorescence anisotropy measurements, both derived from an 80- nucleotide palindromic oligonucleotide comprising the TelN target sequence *ntelR* followed by its reverse complement. A fluorescein (6FAM) label was attached to the 3‘ end of the oligonucleotide: 5’-aattacggaacatatcagcacacaattgcccattatacgcgcgtataatgggcaattgtgtgctgatatgttccgtaatt-[6FAM]-3’

For the 40 bp specific *ntelR* hairpin substrate, the oligonucleotide was subjected to intramolecular annealing in buffer (100 mM NaCl, 10 mM Tris-HCl pH 8, and 1 mM EDTA) by heating to 95°C for 1 minute, followed by rapid cooling on ice. The 80 bp specific *ntelRR* double-stranded DNA was prepared using the same oligonucleotide through intermolecular annealing at 25°C overnight in the same buffer. Fluorescence anisotropy measurements were performed in a reaction buffer containing 25 mM Tris pH 7.5, 50 mM KCl, 5 mM MgCl3, and 1 mM MnCl3. A series of TelN protein concentrations, ranging from 0 to 9.6 μM, were incubated in the presence of 50 nM fluorescein-labeled DNA substrates for 30 min at room temperature to attain equilibrium. Measurements were recorded using a Synergy Neo Hybrid Multi-Mode Microplate reader equipped with appropriate fluorescence polarization filters. Assays were conducted in black 96-well flat-bottom plates at 25°C. Data analysis and curve fitting were performed using GraphPad Prism 10 software using non-linear regression to determine binding parameters.

### Purification of Mre11-Rad50 protein complex

Mre11-Rad50 was purified as described in (Roisné-Hamelin et al., 2024). The Mre11-Rad50 protein complex was expressed in *E. coli* BL21, using a dual vector system (N-terminal 10His-TwinStrep-3C tagged Mre11 and untagged Rad50). Cells were grown in TB medium at 37°C until OD600=1, cooled to 18°C, and protein expression was induced with 0.5 mM IPTG for 16 hours. Cells were harvested and resuspended in lysis buffer (50 mM Tris pH 7.5, 300 mM NaCl, 5% glycerol, 25 mM imidazole) containing 100 μL of protease inhibitor cocktail and 5 mM β-mercaptoethanol. Cells were lysed by sonication on ice, using a VS70T probe mounted on a SonoPuls unit (Bandelin), at 40% output power for 13 min with pulsing (1 s on/1 s off). After sonication and ultracentrifugation (40,000 g, 45 min), the clarified lysate underwent a three-step chromatographic purification. First, metal affinity chromatography using a HisTrap HP 5 mL column with imidazole gradient elution (25-500 mM); second, anion exchange chromatography on a HiTrap Q 5 mL column eluted with a NaCl gradient (50-1000 mM); and finally, a size-exclusion chromatography on a Superose 6 Increase 10/300 GL column equilibrated with 20 mM Tris-HCl pH 7.5, 250 mM NaCl and 1 mM TCEP. The peak fractions containing the intact complex were concentrated to approximately 2.3 mg/mL, flash-frozen in liquid nitrogen, and stored at –70°C.

### Purification of TelN constructs

TelN constructs (TelN WT, TelN Y425F, TelN Δ540-631, TelN Δ582-631) with N-terminal His-TwinStrep- 3C tags were expressed in *E. coli* BL21 cells. Cultures were grown in TB medium at 37°C until OD600=1, cooled to 20°C, and protein expression was induced with 0.5 mM IPTG for 16 hours. Cells were harvested by centrifugation and resuspended in lysis buffer (50 mM Tris pH 7.5, 300 mM NaCl, 5% glycerol, 25 mM imidazole) supplemented with 100 μL of protease inhibitor cocktail and 5 mM β- mercaptoethanol. Following sonication and ultracentrifugation (40,000 g, 45 min), the clarified lysate underwent a four-step purification process. First, proteins were purified by affinity chromatography using a HisTrap HP column with imidazole gradient elution (25-500 mM). Then, for TelN WT, TelN Y425F, and TelN Δ582-631, overnight tag cleavage was performed during dialysis using 20 mM Tris pH 7.5, 100 mM NaCl, and 5 mM β-mercaptoethanol supplemented with 500 μL 3C protease. For TelN Δ540-631, tag cleavage was performed overnight without dialysis to prevent precipitation. Finally, proteins were subjected to anion exchange chromatography using a HiTrapQ column with NaCl gradient elution. Finally, size-exclusion chromatography was performed on a HiLoad Superdex 200 column equilibrated with 10 mM Tris pH 7.5, 200 mM NaCl, 0.1 mM EDTA, and 1 mM DTT. Purified proteins were concentrated to 7.4 mg/mL (TelN WT), 7.0 mg/mL (TelN Y425F), 4.0 mg/mL (TelN Δ582-631), and 4.3 mg/mL (TelN Δ540-631), flash-frozen in liquid nitrogen, and stored at -70°C.

### DNA cleavage assay

37 nM of 566 bp *ntelRL* DNA sequence, generated by PCR amplification, was incubated with increasing concentrations of TelN constructs (0-1200 nM) in 20 μL reactions containing nuclease buffer (25 mM Tris-HCl pH 7.5, 50 mM KCl, 5 mM MgCl3, 1 mM MnCl3, 0.1 mg/mL BSA, 1 mM DTT). Samples were incubated at 30°C for 30 minutes, followed by heat inactivation at 75°C for 5 minutes. The resulting DNA species were resolved by electrophoresis on 1.5% agarose gels containing ethidium bromide.

### DNA end protection assay

A 566 bp *ntelRL* sequence (*ntelRL* cognate sequence with additional neighboring sequences from N15 phage) was incubated with TelN, generating two hairpin products of 237 bp and 329 bp using its cleaving-joining activity. The biotinylated and fluorescently labeled 237 bp hairpin substrate was used for subsequent analyses. DNA immobilization was performed using T1 streptavidin-coated Dynabeads (10 μL, 100 μg per reaction). Beads were first pre-equilibrated using 10 volumes of nuclease buffer (25 mM Tris-HCl, pH 7.5, 50 mM KCl, 5 mM MgCl3, 1 mM MnCl3, 0.1 mg/mL BSA, 1 mM DTT). 37 nM of biotinylated hairpin DNA was bound to the beads in a 20 μL reaction at 25°C for 15 minutes with constant shaking at 850 rpm. Protection assays were conducted by incubating the immobilized hairpin DNA with Mre11-Rad50 tetramer (125 nM) and increasing concentrations of TelN (0-1200 nM). Reactions were performed in nuclease buffer supplemented with 1 mM ATP at 37°C for 15 minutes with shaking. Reactions were stopped by adding 80 μL buffer containing 10 mM EDTA and 20 mM Tris.

Protected DNA was eluted by adding 100 μL pre-heated nuclease buffer supplemented with 25 mM biotin and 0.1% SDS at 70°C for 15 minutes. The supernatant containing the eluate was separated from the beads using a magnetic rack. Samples were resolved on 5% denaturing polyacrylamide gels prepared in 1× alkaline buffer (50 mM NaOH, 1 mM EDTA). Gels were processed under light-protected conditions and subsequently scanned using a Typhoon fluorescence imager (GE Healthcare). Band intensities were quantified using ImageQuant™ TL analysis software.

For species specificity experiments, 1 nM of fluorescently labeled hairpin DNA substrate was incubated with either 80 nM *E. coli* MR or 80 nM *S. cerevisiae* MRX complemented with 320 nM pSae2. Both MRX and pSae2 were kindly provided by the Cejka lab. TelN was added at three different concentrations (20 nM, 80 nM, and 320 nM) to assess protection against both complexes. The reactions were performed and analyzed following the same protocol described above.

For multi-substrate protection assays, we used three different fluorescently labeled DNA substrates simultaneously: a 329 bp *ntel*-containing hairpin labeled with TAMRA, a biotinylated 237 bp *ntel*- containing hairpin labeled with fluorescein and immobilized on streptavidin beads, and a 411 bp non- specific linear DNA fragment (random sequence, non-*ntelRL*, non-hairpin DNA) labeled with ATO 680. The total DNA concentration was maintained at 30 nM across all experiments, with equimolar amounts of each substrate (10 nM each). Protection assays were conducted by incubating the DNA substrates with 500 nM Mre11-Rad50 tetramer and increasing concentrations of TelN WT (0-640 nM). Reactions were performed in nuclease buffer supplemented with 1 mM ATP at 37°C for 15 minutes with shaking, followed by the same stopping and elution procedures described above. For each fluorophore (fluorescein, TAMRA, and ATO 680), the Typhoon fluorescence imager (GE Healthcare) was set with appropriate excitation and emission filter settings. For overlay images, individual fluorescence channels were pseudo-colored (fluorescein: blue, TAMRA: green, and ATO 680: red) and combined using Adobe Photoshop.

### Southern blot analysis

Genomic DNA was extracted from *B. subtilis* strains using standard methods and digested with BsaI and XhoI restriction enzymes (New England Biolabs) overnight at 37°C. Digested DNA was separated on 0.8% agarose gels and transferred to nylon membranes (Hybond-N+, GE Healthcare) using alkaline transfer methods. A 32P-labeled DNA probe targeting the *ntelRL*-spectinomycin resistance gene region was hybridized overnight at 65°C. Membranes were washed with 2× SSC/0.1% SDS at room temperature and 0.1× SSC/0.1% SDS at 65°C, then exposed to a phosphor image screen and subsequently scanned on a Typhoon scanner (GE Healthcare). Fragment sizes were determined using DNA molecular weight standards.

