## Supplemental Figures and Legends for "Phage-Encoded TelN Inhibits Mre11-Rad50 to Protect Hairpin Telomeres"

**A**

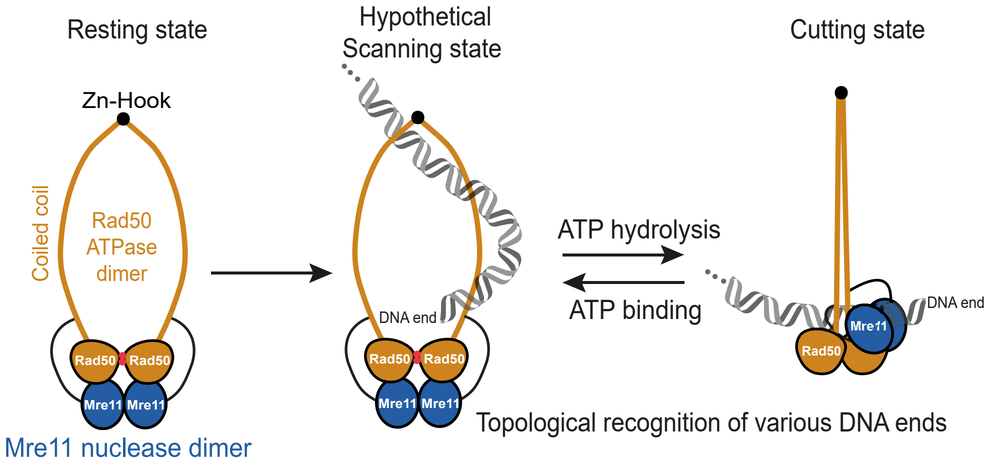


Model for *E. coli* MR sensing and processing of linear DNA ends

**B**

Summary of *B. subtilis* suppressor mutations in the *specR–nteIRL* locus

| **Mutant** | **Mutation Type** | **Location** | **Nucleotide Substitution / Deletion** |
|---|---|---|---|
| Mutant 1 | Substitution | *nteIRL* site | C19T |
| Mutant 2 | Substitution | *nteIRL* site | T15A, C17T, C19A |
| Mutant 3 | Deletion | *specR–nteIRL* locus | 511 bp deletion |
| Mutant 4 | Deletion | *specR–nteIRL* locus | 225 bp deletion |

```
               1            nteIR                           nteIL               56
nteIRL site  TATCAGCACACAATTGCCCATTATACGCGCGTATAATGGACTATTGTGTGCTGATA
mutant 1     TATCAGCACACAATTGCCTATTATACGCGCGTATAATGGACTATTGTGTGCTGATA
mutant 2     TATCAGCACACAATAGTCAATTATACGCGCGTATAATTGACTATTGTGTGCTGATA
```

**C** **D**

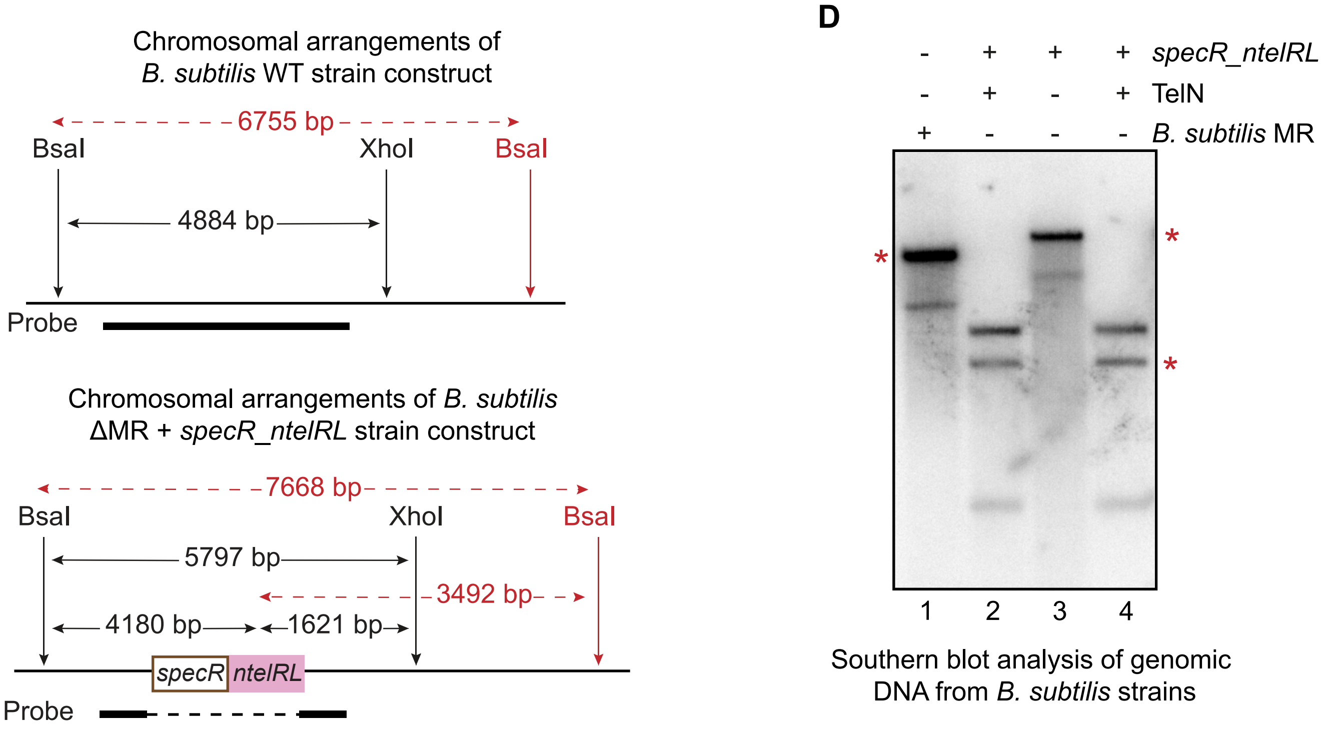


Southern blot analysis of genomic DNA from *B. subtilis* strains

**E**

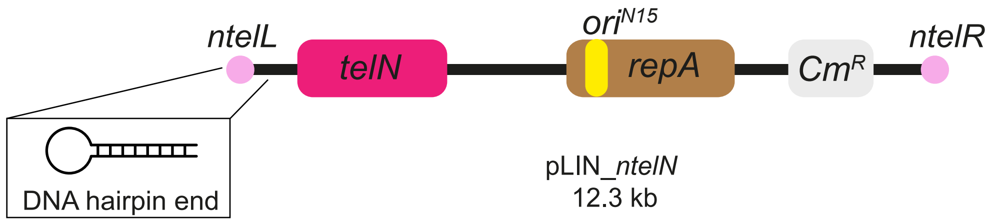


**Figure S1**

**Supplementary figure legends**

**Figure S1. (A)** Schematic of the proposed model for *E. coli* MR sensing and processing of chemically diverse DNA ends. Rad50 comprises an ATP-regulated catalytic head harboring DNA-binding activities, two long, protruding antiparallel coiled coils, and an apical zinc-hook dimerization domain. The DNA-binding and processing head module is formed by the Mre11 nuclease dimer and the Nucleotide-Binding Domains (NBDs) of Rad50. In the resting state, the complex adopts an auto-inhibited conformation where the Rad50 ATPase dimer blocks the Mre11 nuclease active site. Upon DNA binding, the complex transitions to a hypothetical scanning state where it searches for DNA ends along the double helix. Detection of a DNA end triggers an ATP-regulated conformational change, forming the cutting state where a ring-to-rod transition occurs as the two coiled coils close onto a single DNA double helix, repositioning the Mre11 dimer to the side of the complex. MR specifically processes linear DNA termini and discriminates against circular DNA, as the presence of two DNA strands within the ring formed by MR would prevent the closure of the Rad50 coiled coils. **(B)** Suppressor mutations in the *specR–ntelRL* locus in *B. subtilis*. To test whether the *B. subtilis* chromosome can be linearized, we co-introduced *telN* and its cognate *ntelRL* site. No clones carrying both intact *telN* and *ntelRL* were recovered; rare suppressor clones carrying mutations in *ntelRL*. Two-point mutants (mutants 1 and 2) contain substitutions within the *ntelRL* site (red boxes). Mutant 1 carries a single substitution (C19T); Mutant 2 harbors three (T15A, C17T, C19A). Two additional suppressor clones (Mutants 3 and 4) contain large deletions (511 bp and 225 bp, respectively) spanning the *specR–ntelRL* locus. **(C)** Schematic representation of the analyzed chromosomal regions for Southern blot analysis. Genomic DNA was digested with BsaI and XhoI. The upper scheme shows the wild-type (WT) configuration with expected fragments: 4884 bp between BsaI and XhoI (black arrows) and 6755 bp between both BsaI sites in case of partial XhoI digestion (red dashed arrows). The position of the probe used for hybridization is indicated by a black bar targeting the *ntelRL*-spectinomycin region. The lower scheme shows the ΔMR + *specR_ntelRL* construct, characterized by the deletion of *B. subtilis* Mre11-Rad50 with insertion of both the spectinomycin resistance gene and the *ntelRL* sequence (*ntelRL* with additional neighboring sequences from N15 phage). The primary focus is on BsaI-XhoI fragment patterns (black arrows): 5797 bp in the intact construct. Due to partial XhoI digestion observed in our experiments, additional BsaI-BsaI fragments are also shown (red dashed arrows): 7668 bp in the intact construct. When TelN cleaves at the *ntelRL* site, the BsaI-XhoI fragment is split into two fragments of 4180 bp and 1621 bp. In partially XhoI-digested DNA with TelN processing, a 3492 bp fragment would appear between the cleaved *ntelRL* site and the second BsaI site (in red). **(D)** Southern blot analysis of BsaI/XhoI-digested genomic DNA from different *B. subtilis* strains. While our analysis primarily focuses on BsaI-XhoI digested fragments, partial XhoI digestion resulted in additional bands (marked with red asterisks) representing BsaI-BsaI fragments. Lane 1: WT strain showing the expected 4884 bp BsaI-XhoI band and a 6755 bp band (*) resulting from partial XhoI digestion (BsaI-BsaI). Lanes 2, 3, and 4: Strains lacking the *B. subtilis* MR complex (ΔMR) and containing the *ntelRL* site inserted adjacent to the spectinomycin resistance gene (*specR*). Lane 2 (ΔMR + *specR_ntelRL* + $P_{xyl}$-*TelN*) and lane 4 (ΔMR + *specR_ntelRL* + *PrpsB-TelN*) express TelN under the xylose-inducible promoter ($P_{xyl}$) or the constitutive ribosomal protein S2 promoter (PrpsB), respectively. These strains display two distinct bands at 1622 bp and 4180 bp from BsaI-XhoI digestion, confirming that TelN cleaves at the *ntelRL* site, resulting in chromosome linearization. An additional 3492 bp band (*) appears due to partial XhoI digestion in these linearized chromosomes. Lane 3 (ΔMR + *specR_ntelRL* + PrpsB-GFP): Strain containing the *ntelRL* site but expressing GFP instead of TelN shows a 5797 bp BsaI-XhoI band and a 7668 bp band (*) from partial XhoI digestion, demonstrating that chromosome linearization specifically requires TelN expression. **(E)** Schematic of pLIN_*ntelN* linear plasmid (12.3 kb) with its *ntel* DNA hairpin ends (*ntelL* and *ntelR*) and its own copy of the *telN* gene, depicted in the blowout, adapted from (Liu et al., 2022).

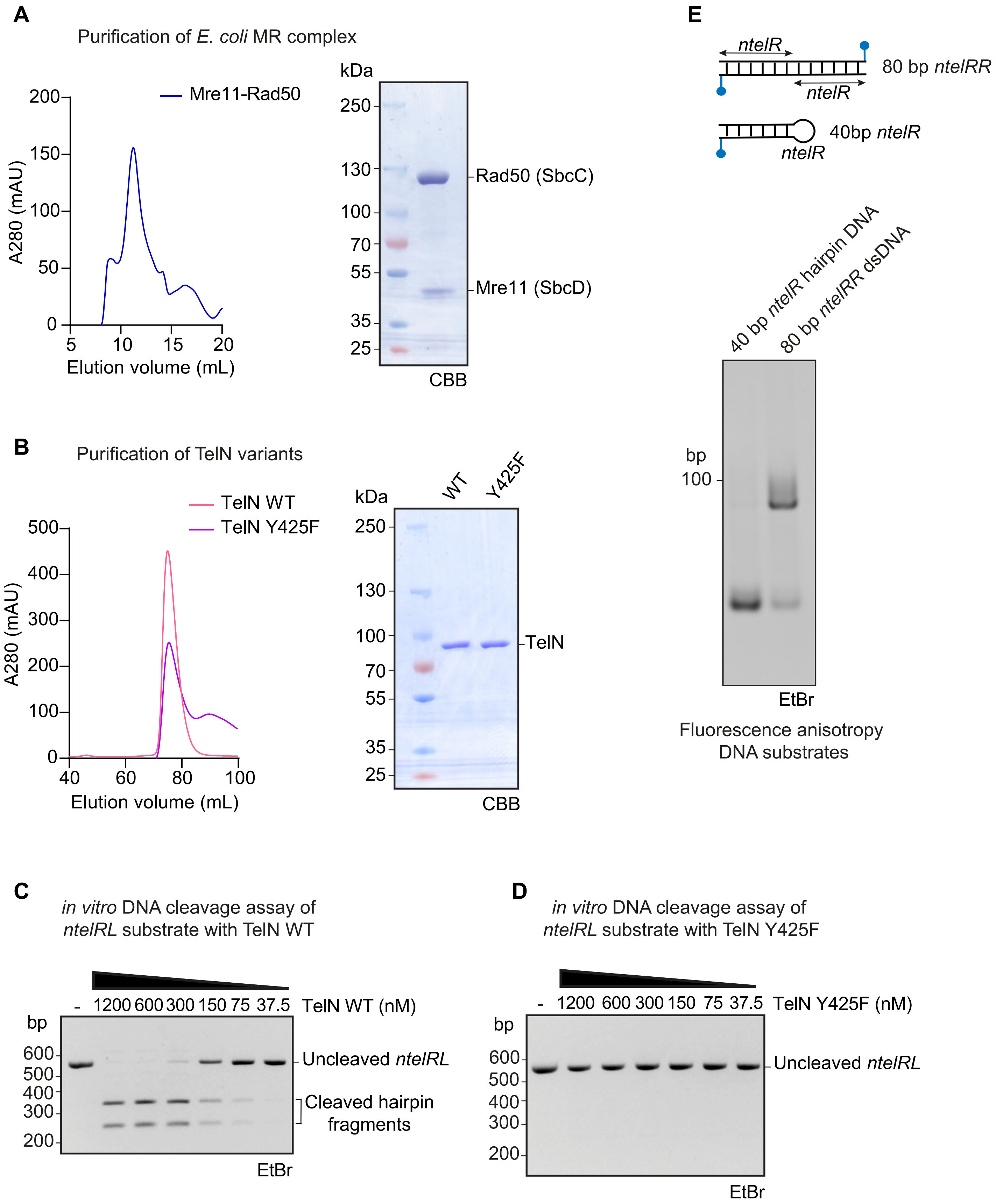
A
Purification of *E. coli* MR complex
Mre11-Rad50
A280 (mAU)
200
150
100
50
0
5
10
15
20
Elution volume (mL)
kDa
250
130
100
70
55
35
25
Rad50 (SbcC)
Mre11 (SbcD)
CBB
B
Purification of TelN variants
TelN WT
TelN Y425F
A280 (mAU)
500
400
300
200
100
0
40
60
80
100
Elution volume (mL)
kDa
WT
Y425F
250
130
100
70
55
35
25
TelN
CBB
C
*in vitro* DNA cleavage assay of
*ntelRL* substrate with TelN WT
- 1200 600 300 150 75 37.5 TelN WT (nM)
bp
600
500
400
300
200
Uncleaved *ntelRL*
Cleaved hairpin
fragments
EtBr
D
*in vitro* DNA cleavage assay of
*ntelRL* substrate with TelN Y425F
- 1200 600 300 150 75 37.5 TelN Y425F (nM)
bp
600
500
400
300
200
Uncleaved *ntelRL*
EtBr
E
*ntelR*
*ntelR*
80 bp *ntelRR*
40bp *ntelR*
*ntelR*
40 bp *ntelR* hairpin DNA
80 bp *ntelRR* dsDNA
bp
100
EtBr
Fluorescence anisotropy
DNA substrates

**Figure S2**

**Figure S2. (A-B)** Left panels: representative size exclusion chromatography (SEC) elution profile for the purification of **(A)** the *E. coli* MR complex, **(B)** TelN WT and TelN Y425F proteins. Right panels: SDS-PAGE profile of (A) purified MR, **(B)** purified TelN WT and TelN Y425F visualized by Coomassie brilliant blue (CBB) staining. Proteins were diluted to 1 µM before loading on a 4-12% gradient acrylamide gel. **(C-D)** DNA cleavage assays on *ntelRL* DNA substrate (*ntelRL* with additional neighboring sequences from N15 phage) using decreasing concentrations of **(C)** TelN WT and **(D)** TelN Y425F. DNA species were resolved on a 1.5% ethidium bromide (EtBr) agarose gel. The uncleaved *ntelRL* linear substrate runs at 566 bp, while TelN cleavage results in two hairpin fragments of 329 bp and 237 bp. **(E)** Analysis of fluorescently labeled DNA substrates on a 10% native polyacrylamide gel. The substrates used for binding measurements with TelN WT and TelN Y425F include a 40 bp *ntelR* hairpin DNA and an 80 bp *ntelRR* double-stranded DNA. Both substrates were derived from the same 80-nucleotide palindromic oligonucleotide, which contains the *ntelR* sequence followed by its reverse complement. The palindromic nature of this sequence allows it to form either a 40 bp hairpin through intramolecular annealing or an 80 bp dsDNA through intermolecular annealing, depending on preparation conditions.

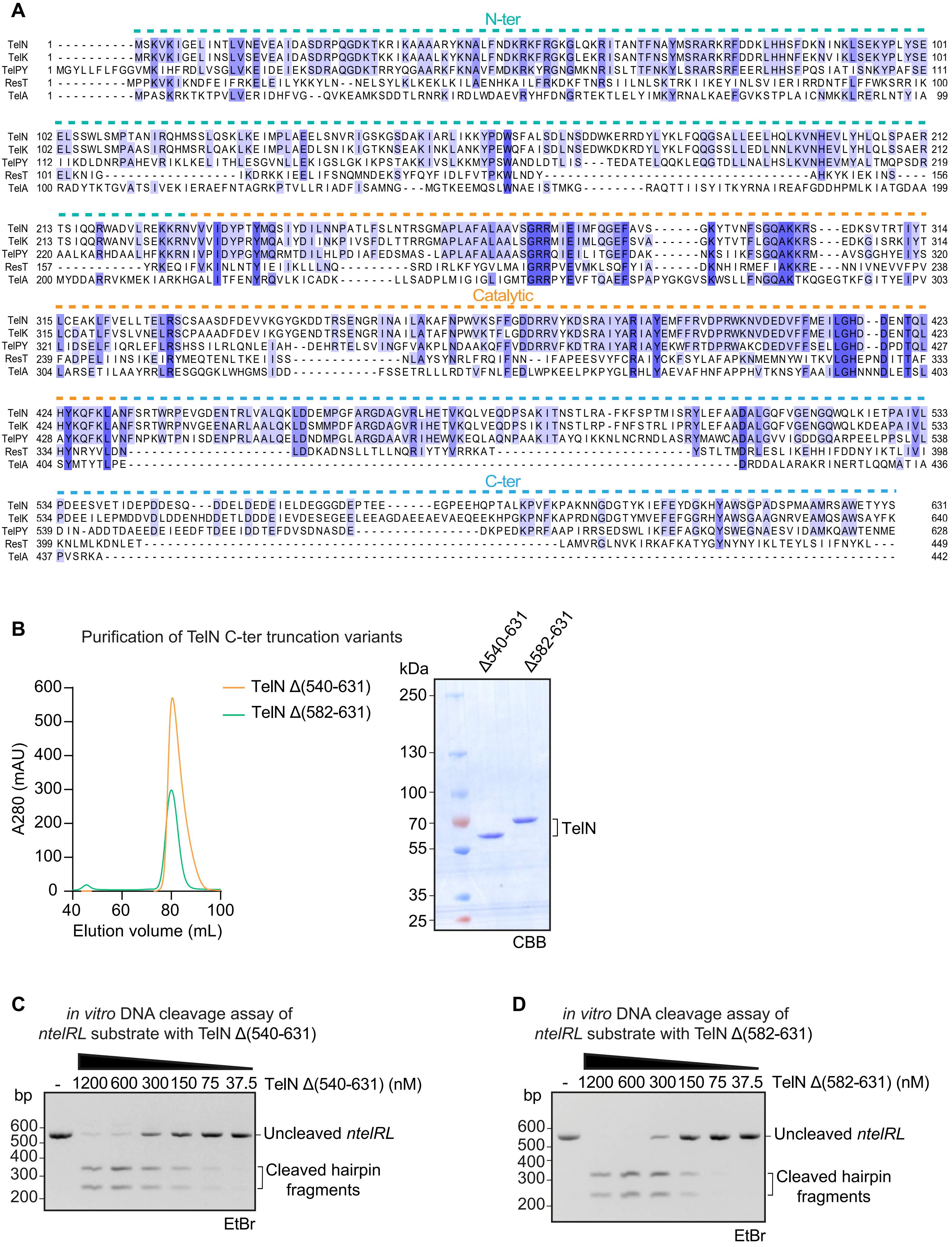

A
N-ter
Catalytic
C-ter
TelN
TelK
TelPY
ResT
TelA
B
Purification of TelN C-ter truncation variants
TelN Δ(540-631)
TelN Δ(582-631)
A280 (mAU)
Elution volume (mL)
kDa
Δ540-631
Δ582-631
TelN
CBB
C
in vitro DNA cleavage assay of ntelRL substrate with TelN Δ(540-631)
bp
- 1200 600 300 150 75 37.5 TelN Δ(540-631) (nM)
Uncleaved ntelRL
Cleaved hairpin fragments
EtBr
D
in vitro DNA cleavage assay of ntelRL substrate with TelN Δ(582-631)
bp
- 1200 600 300 150 75 37.5 TelN Δ(582-631) (nM)
Uncleaved ntelRL
Cleaved hairpin fragments
EtBr


**Figure S3**

**Figure S3. (A)** Multiple-sequence alignment of telomere resolvases from different organisms. Alignment includes TelN from *E. coli* phage N15, TelK from *Klebsiella oxytoca* phage phiKO2, TelPY from *Yersinia* phage PY54, TelA from *Agrobacterium tumefaciens*, and ResT from *Borrelia burgdorferi*. Numbers indicate amino acid positions. The sequence alignment was done on JalView, and the residues are color-coded based on percentage identity. The TelN domains are indicated by colored dashed lines: N-ter domain (green dashed line), catalytic domain (orange dashed line), and C-ter domain (blue dashed line). (Sequence Accession numbers: TelN: Q37967, TelK: Q6UAV6, TelPY: Q7Y3Y3, ResT: O50979, TelA: F8KAE9). **(B)** Left panel: representative size exclusion chromatography (SEC) elution profiles for the purification of TelNΔ540-631 and TelNΔ582-631 truncation proteins. Right panel: SDS-PAGE profiles of purified proteins visualized by Coomassie brilliant blue (CBB) staining. Proteins were diluted to 1 µM before loading on a 4-12% gradient acrylamide gel. **(C-D)** DNA cleavage assays on *ntelRL* DNA substrate using decreasing concentrations of **(C)** TelNΔ540-631 and **(D)** TelNΔ582-631. DNA species were resolved on a 1.5% EtBr agarose gel. The uncleaved *ntelRL* linear substrate runs at 566 bp, while TelN cleavage results in two hairpin fragments of 329 bp and 237 bp.

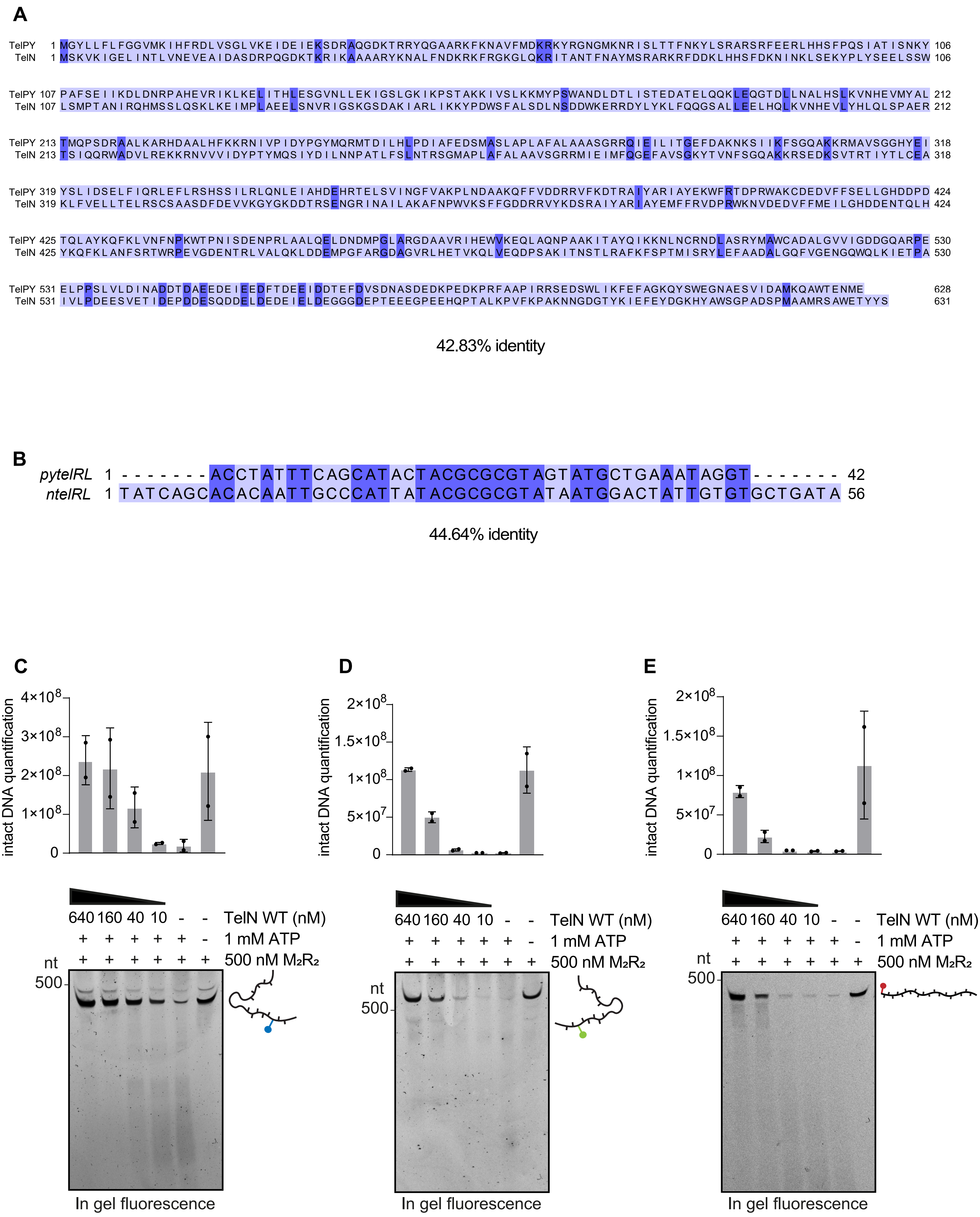
A
TelPY
TelN
42.83% identity
B
pytelRL 1 - - - - - - - ACCTATTTCAGCATACTACGCGCGTAGTATGCTGAAATAGGT - - - - - - - 42
ntelRL 1 TATCAGCACACAATTGCCCATTATACGCGCGTATAATGGACTATTGTGTGCTGATA 56
44.64% identity
C
D
E
intact DNA quantification
640 160 40 10 - -
TelN WT (nM)
1 mM ATP
500 nM M2R2
nt
500
In gel fluorescence

**Figure S4**

**Figure S4. (A)** Sequence alignment of TelPY and TelN proteins. Numbers indicate amino acid positions. The alignment was performed using JalView software, with residues color-coded based on percentage identity. The overall sequence identity is 42.83%, as determined by SnapGene software. (Sequence Accession numbers: TelN: Q37967, TelPY: Q7Y3Y3) **(B)** Sequence alignment of the recognition sequences *pytelRL* (42 bp) and *ntelRL* (56 bp). Numbers above the sequences indicate nucleotide positions. The alignment was performed using JalView software, with nucleotides color-coded based on percentage identity. The overall sequence identity is 44.64%, as determined by SnapGene software. **(C-E)** Individual gel scans corresponding to the overlay image shown in Figure 4C for a clearer interpretation of substrate-specific degradation patterns. Lower panels: representative in-gel fluorescence analysis of DNA products. Upper panels: quantification of intact DNA substrates. The cartoons next to each gel indicate the DNA substrate used in the assay: **(C)** bead-immobilized 237 bp *ntelL* hairpin DNA substrate labeled with blue fluorescence, **(D)** 329 bp *ntelR* hairpin DNA substrate labeled with green fluorescence, **(E)** 411 bp non-specific linear DNA substrate, which lacks the *ntelRL* sequence, labeled with red fluorescence.

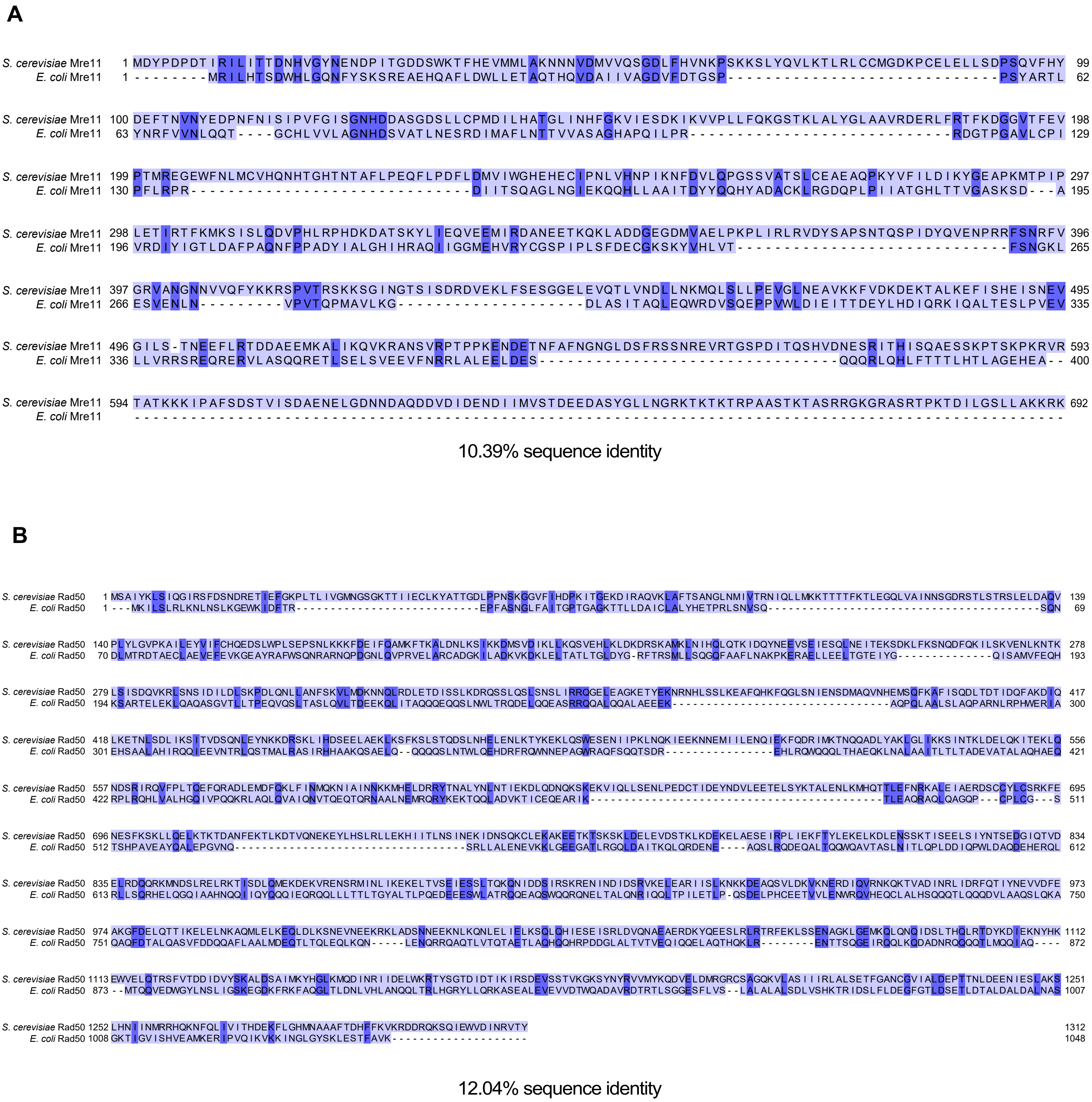
A
S. cerevisiae Mre11
E. coli Mre11
1 99
1 62
100 198
63 129
199 297
130 195
298 396
196 265
397 495
266 335
496 593
336 400
594 692
10.39% sequence identity
B
S. cerevisiae Rad50
E. coli Rad50
1 139
1 69
140 278
70 193
279 417
194 300
418 556
301 421
557 695
422 511
696 834
512 612
835 973
613 750
974 1112
751 872
1113 1251
873 1007
1252 1312
1008 1048
12.04% sequence identity

### Figure S5

**Figure S5. (A)** Sequence alignment of Mre11 proteins from *E. coli* and *S. cerevisiae*. Numbers indicate amino acid positions. The overall sequence identity is 10.39%, as determined by SnapGene software. (Sequence Accession numbers: Mre11: *Saccharomyces cerevisiae*: P32829, *Escherichia coli*: P0AG76). **(B)** Sequence alignment of Rad50 proteins from *E. coli* and *S. cerevisiae*. Numbers indicate amino acid positions. The overall sequence identity is 12.04%, as determined by SnapGene software. (Sequence Accession numbers: Rad50: *Saccharomyces cerevisiae*: P12753, *Escherichia coli*: P13458). Alignments were performed using JalView software, with residues color-coded based on percentage identity.
